# A causality-inspired feature selection method for cancer imbalanced high-dimensional data

**DOI:** 10.1101/2021.10.04.462984

**Authors:** Yijun Liu, Qiang Huang, Huiyan Sun, Yi Chang

**Author notes:** Corresponding author. (HS), (YC).

## Abstract

It is significant but challenging to explore a subset of robust biomarkers to distinguish cancer from normal samples on high-dimensional imbalanced cancer biological omics data. Although many feature selection methods addressing high dimensionality and class imbalance have been proposed, they rarely pay attention to the fact that most classes will dominate the final decision-making when the dataset is imbalanced, leading to instability when it expands downstream tasks. Because of causality invariance, causal relationship inference is considered an effective way to improve machine learning performance and stability. This paper proposes a Causality-inspired Least Angle Nonlinear Distributed (CLAND) feature selection method, consisting of two branches with a class-wised branch and a sample-wised branch representing two deconfounder strategies, respectively. We compared the performance of CLAND with other advanced feature selection methods in transcriptional data of six cancer types with different imbalance ratios. The genes selected by CLAND have superior accuracy, stability, and generalization in the downstream classification tasks, indicating potential causality for identifying cancer samples. Furthermore, these genes have also been demonstrated to play an essential role in cancer initiation and progression through reviewing the literature.

**Author Summary:** Selecting trustworthy biomarkers from high-dimensional data is an important step to help researchers and clinicians understand which genes play key roles in cancer development and progression. A large number of machine learning-based feature selection algorithms have been generated in recent years for biomarker discovery. However, these methods usually show unstable results in the face of class-imbalanced biological data, making it seem unreliable for researchers. Here we introduce the causal theory with the property of causal invariance to aid in the design of feature selection algorithms, analyze how imbalanced distributions affect feature selection methods, and propose a novel causality-based feature selection method. The method with bilateral structure adjusts the data distribution from both class-wise and sample-wise to eliminate the effect of imbalance on the results. Additionally, CLAND can simultaneously address the nonlinearity and high-dimensionality of cancer data, which broaden its application scope. We conducted extensive experiments on six real imbalance cancer datasets and obtained efficient and stable results, while the obtained biomarker has significant biological significance.

## 1 Introduction

Identifying biomarkers with distinguishing ability is a critical step towards cancer diagnosis and prognosis prediction and helps further understand the mechanism of cancer initiation and various phenotypes. Over the years, many computational feature selection methods have been proposed to identify critical biomarkers for cancer and cancer subtypes from the data generated by high-throughput technologies [1]. However, as real data are usually imbalanced for each class, such as a large number of cancer samples versus very few normal samples, the selected features are highly partial to large class [2], and such a subset of features is often worthless even if it can achieve high classification accuracy. In addition, most existing feature selection methods have poor robustness and stability when the sample sizes are imbalanced. For the same dataset, features selected by various methods are usually very different. Furthermore, when combining selected features with downstream classification or clustering methods, their performances always vary greatly. The instability raises serious doubts about the reliability of the selected genes as candidate biomarkers [3].

Conventionally, this issue is attributed to the improper assumption of relatively balanced distribution among different classes, and researchers have put forward a series of methods to address it. The sampling-based method is one of the simplest and most effective types. They use the known dataset distribution to re-balance the data distribution, including undersampling for majority class and oversampling for minority, thereby strengthening the learning of the minority [4]. Nevertheless, this method destroys the original data distribution, leading to over-fitting to the minority class or under-fitting to the majority. Another popular type is the cost-sensitive learning method based on heuristic strategies. They add some constraints to weight conditions based on the original standard loss function so that the calculation of the final loss is partial to a specific direction to reduce the bias to the majority class. However, such methods usually require appropriate prior knowledge to establish a corresponding cost matrix [5].

However, regarding the poor performance of traditional feature selection methods for imbalanced data, we suggest the fundamental reason is that the features are obtained by the association with sample labels but not the stable causal relationship. Unlike the association, causality is invariable and can always be identified no matter the data distribution. It has been widely used in economics [6] and epidemiology[7] for many years. Moreover, the introduction of causal mechanisms in the machine learning methods has been demonstrated to enhance their performance, stability, and interpretability [8-10]. Hence, causal inference has attracted more and more attention and has been applied in many scenarios, including image classification and recognition, reinforcement learning and transfer learning. The biggest challenge for causal inferences from observed data is to remove confounders, which are the common causes of the treatment variable and outcome. Assuming imbalanced distribution as the main confounder of selected features and sample labels prediction to identify cancer genes through re-examining and solving the problems of imbalanced transcriptomic data, we propose a novel feature selection method based on causal invariance called Causality-inspired Least Angle Nonlinear Distributed (CLAND).

We design a two-branch structure representing two deconfounder strategies respectively to remove the influence of imbalanced data distribution for feature selection. This structure can prevent the overfitting of the minority class without losing data information. Combining with Hilbert–Schmidt Independence Criterion Lasso, CLAND can simultaneously address other issues of biological cancer data, such as the extremely high-dimensional and non-linear association between features and sample labels. When applying CLAND into several sets of imbalanced cancer transcriptomic data, the selected features can distinguish between cancer and normal samples well and outperform state-of-the-art methods on efficiency and stability. Additionally, several biomarkers obtained by our method have considerable biological significance and have been widely recognized in clinical trials and cancer treatment.

## 2 Related works

### Feature selection methods

The methods can be divided into filter, wrapper, and embedded [11, 12]. Specifically, embedded methods embed feature selection into the process of model construction. As the relationships between biological factors are usually non-linear, we need a feature selection algorithm for high-dimensional data to capture the non-linear relationship between input and output. Minimum redundancy and maximum relevance (mRMR) [13] is a widely used non-linear feature selection method, which uses mutual information as the evaluation measure and selects the most relevant features to the output and is independent of the others. Efficient and robust feature selection (RFS) [14] reduces the influence of noise data by using *l*_1,2_ norm in the loss function and regularization and obtains sparse feature groups simultaneously. Hilbert–Schmidt Independence Criterion Lasso (HSIC Lasso) [15], which can find a small number of features from high-dimensional data in a non-linear manner, can be considered a convex variant of mRMR, a state-of-the-art method of non-linear feature selection. However, these methods do not consider the imbalance of data sets.

### Re-balancing training

The core idea of the most widely used solution to the imbalance problem can be said to re-balance the contribution of different classes in the training phase. It can be divided into three categories: data-level method, algorithm-level method, ensemble method. The data-level method mainly modifies the number of samples in the dataset to make it suitable for standard learning, including under-sampling approaches [16, 17], over-sampling approaches [18, 19], and hybrid approaches [20, 21]. The algorithm-level method mainly modifies the existing methods to reduce the tendency of majority classes. Cost-sensitive learning[22, 23] is the commonly used strategy. The ensemble method combines a data-level or algorithm-level method with an ensemble learning method to obtain a robust strong classifier. However, these integration-based methods are sensitive to noise and have poor applicability.

### Casual inference

It has been demonstrated that machine learning methods could improve their interpretability, transferability, and stability when integrating causal invariance [24]. For example, image classification and detection task in computer vision: [25] uses causal inference to eliminate prediction bias caused by momentum in the training process; target detection task:[26] uses causal inference to eliminate spurious associations between targets and between targets and scenes. In addition to the field of computer vision, the research of causal inference-assisted machine learning also focuses on learning-to-rank[27, 28] and recommendation [29, 30], etc, which apply the user’s implicit feedback as the label.

## 3 Methodology

### Symbol definition

Let *X* = (*x*_1_, …, *x*_*n*_)^*T*^ = (*u*_1_, …, *u*_*m*_) ∈ i ^*n*×*m*^ denote the input data with n samples with m features, and *Y* = (*y*_1_,…, *y*_*n*_)^*T*^ ∈ i ^*n*^ denote the output (or labels) in which *y*_*i*_ ∈*Y* is the label of x_i_. This paper only considers feature selection for the binary classification problem, in which the class with fewer samples is minority class also marked as positive class P and the class with more samples is majority class also marked as negative class N, and use imbalance ratio (IR) to quantify the degree of class imbalance of a dataset as follow:

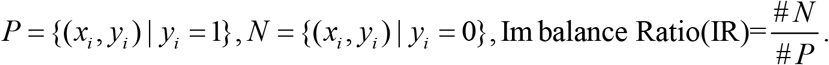

The goal of feature selection from biological data is to select k (k <<m) features most relevant to the label by exploiting the biological data 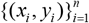.

### Evaluation criteria

In this article, we use the feature selection method to selected the task-related features from the dataset, but we cannot directly evaluate the effect of the selected features. Therefore, we evaluate the amount of information retained in the subset feature by evaluating the performance of different classifiers based on these features.

#### 3.1 A Casual view on the class distribution

To systematically study how imbalanced class distribution affects feature selection, we introduce confounder [31, 32], the common cause of treatment and outcome, and the main factor leading to spurious statistical correlation. Deconfounder can ensure the stability of learning to a certain extent. For instance, considering the relationship between “yellow finger” and “lung cancer,” it is not difficult to find that many people with yellow fingers are more likely to develop lung cancer. Nevertheless, we cannot say that yellow fingers can cause lung cancer, and obviously, there is no causal relationship between them. As we know that smokers are prone to lung cancer, and smokers are also prone to yellow fingers, that is, “yellow finger”←smoke→”lung cancer.” Smoke is the common cause of infection and death and makes a “pseudo correlation” between infection and death, also called “bias.” Therefore, the causality can be obtained only when the observation data is used correctly, and the influence of confounder (also known as confounder bias) is removed.

Then we constructed a causal diagram [32] in Figure 2 (left), where nodes represent variables and arrows represent the direct causal effect with three variables: label probability distribution (D); the feature subset by a given feature selection method (F); the efficiency of classifiers (E). A causal graph is a directed acyclic graph used to show how the variables interact with each other through causal relationships. D is a confounder in the diagram, which is the common cause of F (via D→F) and E (via D→E).

D→F indicates that the feature subset is selected from the labeled dataset by the feature selection method. D→E, it is evident that the label probability distribution will affect the efficiency of the classifier. Therefore, when we evaluate the subset feature’s information by the prediction efficiency of classifiers (F→E), D is a confounder leading to the confounding association flow from F to E.

Let us make some formal explanations and use the Bayes rule to express correlations in Figure 2 (left):

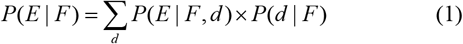

Where each d is the stratification of D and constitutes the whole D. Confounder D introduces the bias through P(d|F). To illustrate, suppose M represents a feature selection method (e.g., mRMR), f* represents the feature set filtered by M, F={f*,M} represents the overall information. For example, P(d=positive-class|F={f*,M}) is small while P(d=negative-class|F={f*,M}) is large. According to Equation 1, the likelihood sum will be attributed to P(E|F={f*,M},d=negative-class) more than P(E|F={f*,M},d=positive-class), so the prediction from F to E will be focused on negative-class rather than the F itself.

To eliminate the influence of imbalanced class distribution (confounder), we adjust D towards balancing label distribution which means D and F is independent. As shown in Figure 2 (right), we balanced the distribution of the label and eliminate the confounding association. Based on the causal view of the influence of label distribution on feature selection, we propose a causality inspired feature selection method called CLAND, which contains three modules and form an efficient and stable feature selection strategy. Specifically, we designed two branches as shown in Figure 1, the class-wise branch and the sample-wise branch and used the ensemble strategy to gather the two branches.

**Figure 1:**
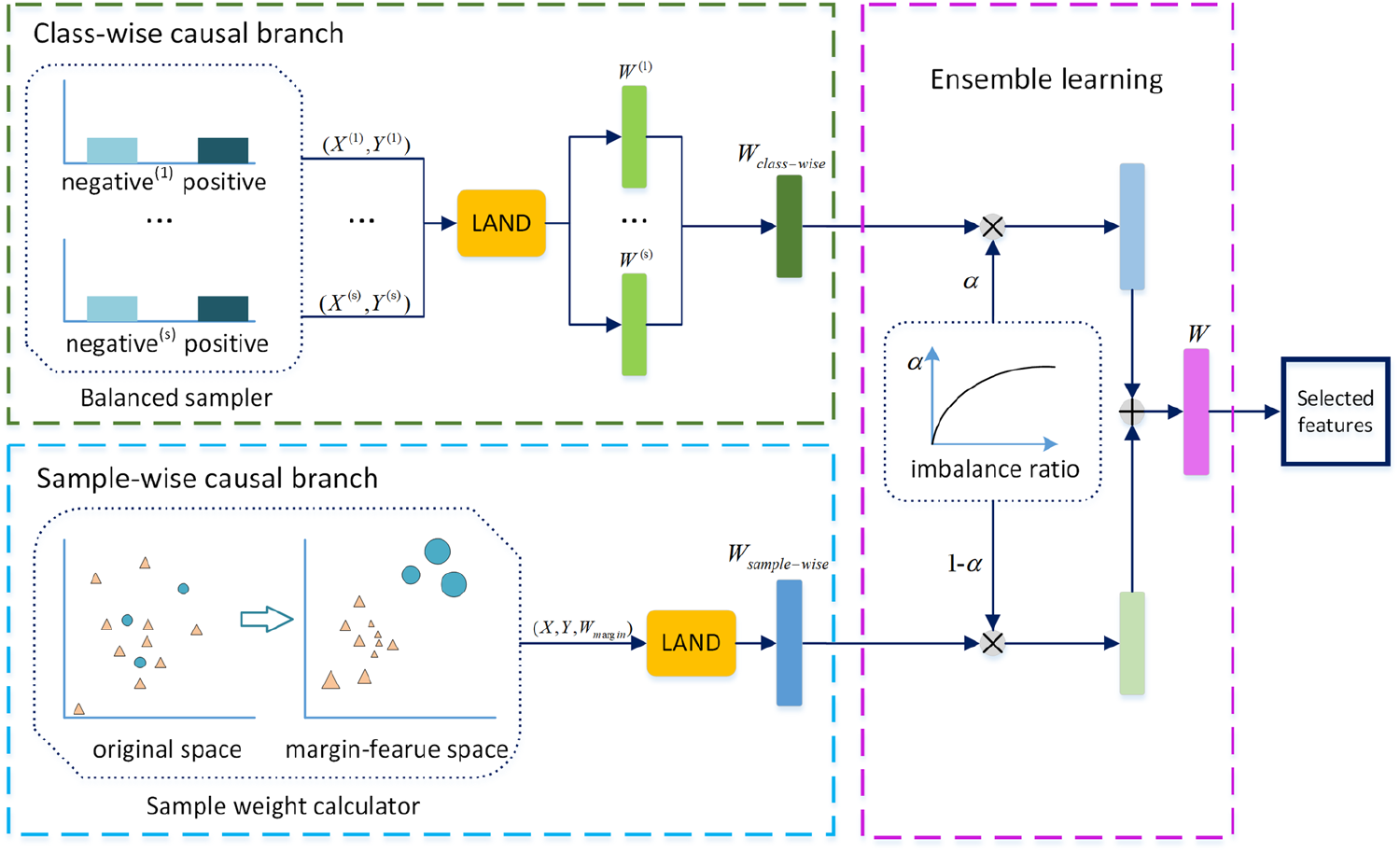
The framework of CLAND consists of three elements: 1) The class-wise causal branch taking re-balanced data as input; 2) The sample-wise causal branch taking the whole data as input; 3) The ensemble learning strategy balances the weights of the features generated by the two branches by using the super parameter *α*.

**Figure 2.**
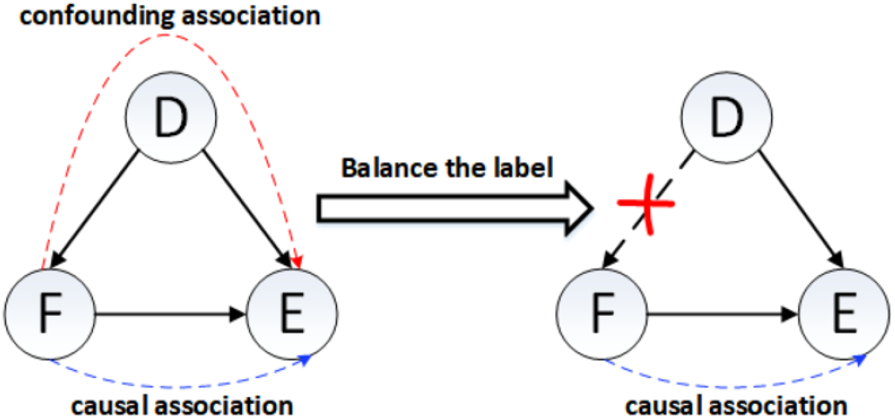
The proposed causal graph. D: the label probability distribution, F: selected features by a given feature selection method, E: efficiency of classifiers.

#### 3.2 Feature selection by LAND

We consider a non-linear extension of LARS [33] leveraging Hibert–Schmidt independence criterion (HSIC) [34] called Least Angle Non-linear Distributed (LAND) feature selection [35].

Specifically, the LAND is a LARS variant of HSIC lasso[15], which can be used to process tens of thousands of features and tens of thousands of samples, and has good prediction ability and interpretability. The optimization problem of HSIC Lasso for paired data 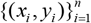 is formulated as:

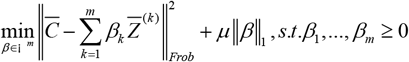

Where ‖g‖_*Frob*_ is the Frobenius norm, ‖g‖_1_ is the *l*_1_ norm, 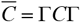 and 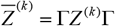 are the centered Gram matrices, 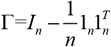 is the centering matrix, *I*_*n*_ is n-dimensional identity matrix, 1_*n*_ is the n-dimensional vector whose elements is all ones. *C*_*ij*_ = *C*(*y*_*i*_, *y* _*j*_) is the delta kernel function for output, 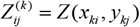 is the Gaussian kernel matrix of the k-th feature input, and *i, j* ∈{1,…, *n*}, *β* = (*β*_1_,…,*β*_*m*_)^*T*^ ∈ i ^*m*^ is the regression coefficient vector, *μ*≥ 0 is the regularization parameter. LAND uses LARS to solve the problem and selects features one by one, and finally gets the most relevant and least redundant feature set. To illustrate the reason. The first term of the objective function can be rewritten as:

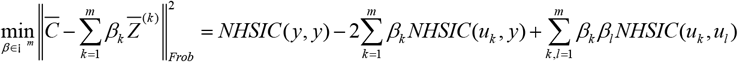

NHSIC(u,y) is the normalized version of HSIC[34] based on kernel function to estimate the dependency between two variables. NHSIC (y,y) is a constant variable and can be ignored in the training process. The larger the value *NHISC*(*u, y*) ∈[0,1], the stronger the dependency between the two variables. If and only if *NHISC*(*u, y*)=0, the two variables are independent and when u=y, *NHISC*(*u, y*)=1. If the dependency between the k-th feature and y is strong, the value of *NHISC*(*u*_*k*_, *y*) is close to 1 and leads to a large value of the regression coefficient β_k_, which means the k-th feature should be preferred. If the k-th feature is independent of y, the value of *NHISC*(*u*_*k*_, *y*) is close to zero, and β_k_ will be very small under the influence of *l*_1_ norm, which means the k-th would not be selected. Moreover, if there is a strong dependency between k-th and l-th features, which means they are redundant with each other,and the value of *NHISC*(*u*_*k*_, *u*_*l*_) is close to one and either of β_k_ and β_l_ would be very small, which ensures that features with the redundant relationship will not be selected at the same time.

LAND iteratively selects non-redundant features strongly related to the output and defines the selection score of the k-th feature as *w*_*k*_ = *NHISC*(*u*_*k*_, *y*) − ∑*β*_*k*\*_ *NHSIC*(*u*_*k*_, *u*_*k*\*_), where k^*^ is the feature that has been selected. Intuitively, this score represents a compromise between the relevance of k-th feature k and output and the degree of redundancy between k-th feature and previously selected features. At the same time, due to the use of HSIC [34], which can capture the non-linear relationship between features and between features and output, the problem of feature-wise non-linear has been solved simultaneously.

#### 3.3 Double branch structure and ensemble strategy

This section, we will explain the two-branch structure proposed in framework and ensemble strategy shown in Figure 1 in detail.

As analysis in Section 3.1, the label probability distribution becomes a confounder while the dataset loss the label balance (IR>1). So for the imbalanced label distribution, we directly adjust it to a balanced distribution in the training process which forces the causal effect from F to E not influenced by imbalance distribution, by the class-wise method and the sample-wise method.

##### Class-wise causal branch

For confounder D, this branch trans it to several balanced datasets *D*′ = {*d* ^(1)^, …, *d* ^(*s*)^} by a balanced sampler and the class-wise implementation is defined as:

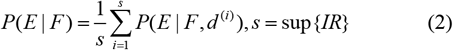

The balanced sampler takes oversampling on the positive class and full sampling on the negative class to use the information contained in all samples fully. Moreover, noise-based data enhancement is set to prevent overfitting. By random sampling without putting back, the negative class is divided into s=sup{IR} sets, and each of them combines with the positive class samples as sub-dataset, and the i-th sub-dataset is labeled as *d* ^(*i*)^ = (*X* ^(*i*)^, *Y* ^(*i*)^). LAND is used to calculate each sub-datasets the feature selection score vector and the i-th weight vector is labeled as *W* ^(*i*)^ ∈ i ^*m*^. Combine s weight vector as the branch weight score by: 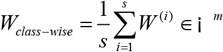.

##### Sample-wise causal branch

This branch is sample-wise, builds a reweighted version of observed distribution by calculating the weight of each sample 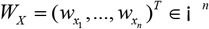 to eliminate the influence of confounder and the implementation is defined as:

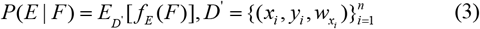

where D^’^ is the dataset with the weight of each sample and f_E_(.) represents the evaluation efficiency of the classifiers. By assigning more weights for positive samples and less for negative samples, the feature selection method would more focus on the positive class in the training process and balance the label distribution on the sample-wise.

To evaluate the affect/weight of each sample on the decision of feature selection, we employ the concept of Margin Vector Feature Space [3] and map the samples in the original feature space to the margin vector feature space by decomposing the margin of a sample along each dimension. For each sample 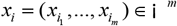 in dataset 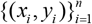 is mapping as 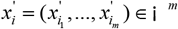, the jth component is formulated as:

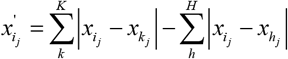

where x_h_ and x_k_ represent the samples with the same and opposite label to x_i_ with the amount K and H, respectively. For the first item in the equation, the value of the positive class with K>H is larger than the value of the negative class with K<H through accumulation operation. And for the second item, the value of the positive class is less than the negative class. So in the new feature space, the positive class has been mapping as a group far away from the negative class, and the degree of deviation increases with the increase of IR.

After the margin vector feature space is generated, the samples in the original space are weighted by the difference of samples in the new space. As the positive class samples always exhibit largely distinct margin vectors from the negative, we assign weights to a sample according to its deviation with rest of samples to increase the weight of the positive class. The formulation of the weight of (*x*_*i*_, *y*_*i*_) we proposed is:

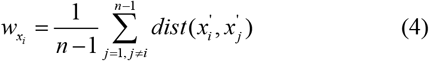

where 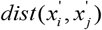 is the Euclidean Distance of two samples in the new feature space.

Therefore, we use the sample-wise causal branch to eliminate the influence of imbalance, and calculate the feature score vector W_sample-wise_ through LAND. We use this branch as a fine grained supplement to the class-wise causal branch.

### Ensemble learning strategy

Integrate the score vectors generated by the two branches to eliminate the influence of confounder from both class-wise and sample-wise. Here, we define a propensity parameter labeled as α and calculated by:

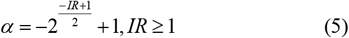

Specifically, the W_class-wise_ is multiplied by the α, the W_sample-wise_ is multiplied by (1-α), and the two new score vectors are added (Figure 1). Furthermore, the feature set is obtained according to the finally obtained score vector W. Although class-wise learning and sample-wise learning are both worthy of attention, as the imbalance ratio increases, our learning focus should shift to the class-wise branch to improve the accuracy of positive class recognition. Therefore, we designed a α-adaptive strategy based on IR. For different data sets, the larger the IR, the larger the α.

## 4 Experiment & Analysis

In this section, we will introduce our experimental results on six real biological datasets. We tested the stability and accuracy of our proposed algorithm to extend to different kinds of classifiers. We also analyzed the effectiveness of the biomarker found by our algorithm. We evaluated the experimental results with multiple criteria and proved the power of our proposed method.

### 4.1 Data source and setup detail

To evaluate the efficiency of our method, we download the transcriptomic data of a total of 2028 samples consisting of 1827 cancer samples and 201 normal samples across six cancer types with different imbalance ratios (IR) from The Cancer Genome Atlas (TCGA) [36] database. We preprocessed datasets and deleted pseudogenes and the genes whose average expression values were less than 10. Table 1 lists the information of each dataset.

**Table 1.** Description of datasets.

| Dataset | #Genes | #Samples | IR |
| --- | --- | --- | --- |
| Kidney Chromophobe (KICH) | 15608 | 91 | 2.64:1 |
| Colon adenocarcinoma (COAD) | 15418 | 311 | 6.59:1 |
| Thyroid carcinoma (THCA) | 15362 | 560 | 8.49:1 |
| Head and Neck squamous cell carcinoma (HNSC) | 15909 | 488 | 10.09:1 |
| Esophageal carcinoma (ESCA) | 5946 | 175 | 12.46:1 |
| Bladder Urothelial Carcinoma (BLCA) | 16088 | 403 | 20.21:1 |

We compared the performance of our proposed method with ReliefF[37], mRMR[13], MIFS[38], REFS[14], LAND[35]. As mentioned in Section 3.1, our proposed method is expected to have a stable performance regardless of the downstream classifier by eliminating the confounders. Therefore, we introduced six classical classifiers to test the effectiveness and stability of different feature selection methods, including NB (Naive Bayes), KNN (K-Nearest Neighbors), LR (Logistic Regression), RF (Random Forest), and GDBT (Gradient Boosting Decision Tree). To better evaluate the models’ performance on imbalanced datasets, we used the confusion matrix as the evaluation index, which can reflect the number of each class that is correctly or incorrectly identified, including AUCPRC[39], F1-score[40], G-mean[41], and MCC[42]. Among them, MCC is the most sensitive to the results of imbalanced datasets.

As the number of features in each dataset is much higher than the sample size (shown in Table 1), all the samples are used for feature selection, and 80% and 20% of them are used as the training set and test set for classifiers, respectively. We use the average result of 10 independent experiments to reduce randomness and show the mean and standard deviation in Table 2 and Supplementary A.1. We expect to retain as much information as possible with fewer features, so the ten most relevant features were selected in each method.

**Table 2.** Detail results of THCA.

| THCA |  |  |  |  |  |  |  |
| --- | --- | --- | --- | --- | --- | --- | --- |
| Model | Metric | ReliefF | mRMR | MIFS | RFS | LAND | CLAND |
| NB | AUCPRC | 0.97±0.008 | 0.889±0.025 | 0.892±0.023 | 0.962±0.015 | 0.927±0.035 | 0.993±0.003 |
|  | F1 | 0.943±0.012 | 0.943±0.012 | 0.943±0.012 | 0.943±0.012 | 0.943±0.012 | 0.969±0.014 |
|  | GM | 0.944±0.011 | 0.944±0.011 | 0.944±0.011 | 0.944±0.011 | 0.944±0.011 | 0.969±0.013 |
|  | MCC | 0.479±0.048 | -0.093±0.15 | -0.044±0.125 | 0.462±0.042 | 0.269±0.172 | 0.795±0.066 |
| KNN | AUCPRC | 0.96±0.014 | 0.917±0.016 | 0.895±0.021 | 0.943±0.022 | 0.993±0.007 | 0.989±0.005 |
|  | F1 | 0.959±0.012 | 0.95±0.01 | 0.943±0.012 | 0.96±0.012 | 0.989±0.005 | 0.985±0.005 |
|  | GM | 0.959±0.012 | 0.951±0.009 | 0.944±0.011 | 0.96±0.012 | 0.989±0.005 | 0.985±0.005 |
|  | MCC | 0.635±0.084 | 0.384±0.1 | 0.051±0.088 | 0.581±0.087 | 0.905±0.039 | 0.868±0.032 |
| LR | AUCPRC | 0.938±0.019 | 0.891±0.021 | 0.892±0.021 | 0.917±0.022 | 0.944±0.027 | 0.992±0.007 |
|  | F1 | 0.956±0.008 | 0.943±0.012 | 0.943±0.012 | 0.946±0.011 | 0.943±0.012 | 0.988±0.004 |
|  | GM | 0.957±0.008 | 0.944±0.011 | 0.944±0.011 | 0.947±0.011 | 0.944±0.011 | 0.988±0.004 |
|  | MCC | 0.524±0.103 | -0.021±0.018 | -0.01±0.016 | 0.334±0.082 | 0.41±0.116 | 0.891±0.046 |
| RF | AUCPRC | 0.97±0.006 | 0.938±0.022 | 0.893±0.02 | 0.966±0.013 | 0.991±0.01 | 0.989±0.009 |
|  | F1 | 0.977±0.005 | 0.954±0.01 | 0.943±0.012 | 0.972±0.01 | 0.989±0.007 | 0.991±0.005 |
|  | GM | 0.977±0.005 | 0.954±0.009 | 0.944±0.011 | 0.972±0.01 | 0.989±0.007 | 0.991±0.005 |
|  | MCC | 0.778±0.04 | 0.502±0.129 | 0.013±0.083 | 0.73±0.094 | 0.904±0.058 | 0.912±0.059 |
| GDBT | AUCPRC | 0.974±0.007 | 0.932±0.021 | 0.894±0.022 | 0.957±0.019 | 0.989±0.008 | 0.989±0.006 |
|  | F1 | 0.982±0.007 | 0.949±0.01 | 0.943±0.012 | 0.968±0.012 | 0.988±0.006 | 0.991±0.005 |
|  | GM | 0.982±0.007 | 0.949±0.01 | 0.944±0.011 | 0.968±0.012 | 0.988±0.006 | 0.991±0.005 |
|  | MCC | 0.828±0.047 | 0.439±0.104 | 0.036±0.071 | 0.679±0.114 | 0.89±0.055 | 0.915±0.039 |

### 4.2 CLAND is comparable to the state-of-the-art methods and has better stability

We use five classifiers with four evaluation criteria to assess the performance of feature selection methods on six imbalanced datasets with different IR. As shown in Figure 3 (a-d), CLAND is superior to other methods in almost all the settings, and the advantages are gradually apparent as the IR of the data set increases. The performance of mRMR, MIFS, and RFS in the KICH dataset with the smallest IR is like other methods, but as the IR increases further, their ability to predict the positive class decreases. When the IR value reached eight or more, the performance of ReliefF, mRMR, MIFS, and RFS all dropped significantly. Moreover, as IR changes, the performance of ReliefF is relatively stable, but it is still lower than LAND and CLAND. Table 2 shows the results of THCA, and the detailed results of other datasets are shown in Supplementary A.1. LAND is the basis of the CLAND method, and it can obtain a good feature set by capturing non-linear relationships between features and labels. However, when it comes to imbalanced data, its efficiency in all data sets is still inferior to CLAND, although higher than other baselines. With the change of IR, it exhibits noticeable oscillation (Supplementary A.1).

**Figure 3:**
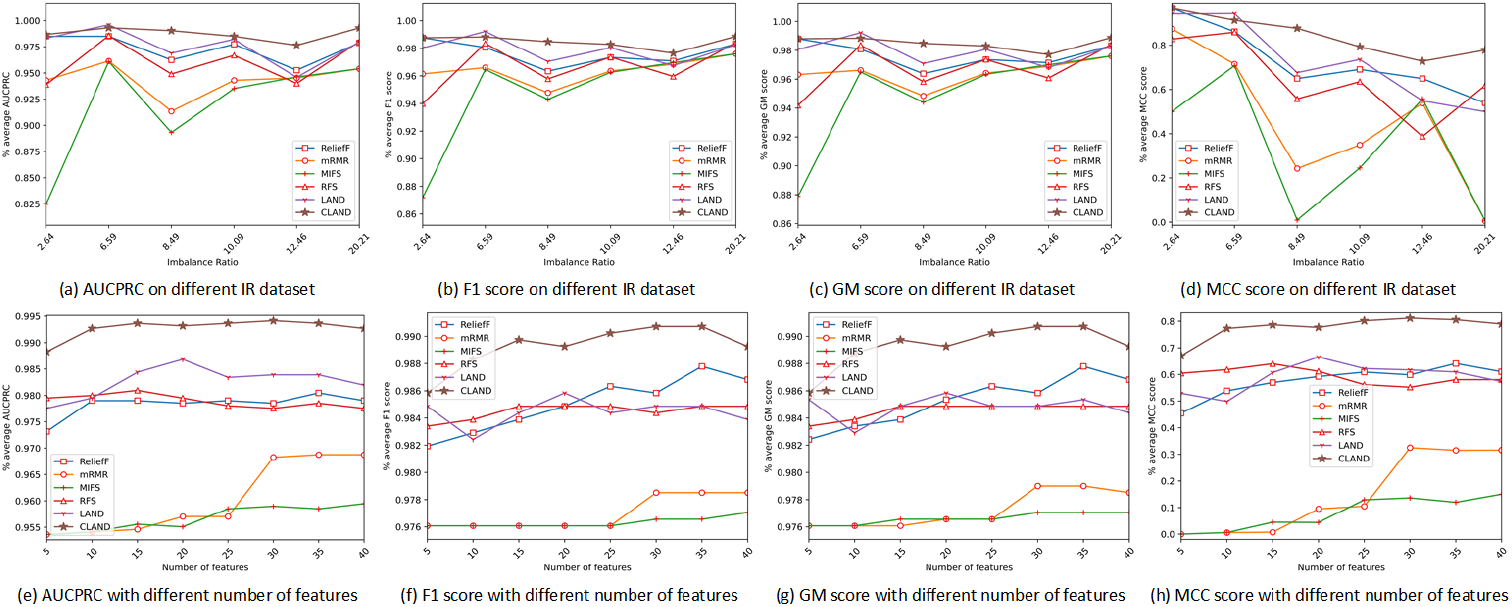
AUCPRC, F1-score, GM, MCC of 6 feature selection methods. The horizontal axis denotes the IR of the dataset in (a, b, c, d) and the number of selected features in (e, f, g, h). The vertical axis denotes the mean value of evaluation metrics.

Figure 3 (a-d) shows CLAND outperforms the traditional feature selection methods in terms of stability. AUCPRC, F1-score, and GM of CLAND all maintained high levels with the increase of IR. Besides, we also evaluate the performance of different selected features numbers on the BLCA dataset and find the performance of CLAND is stable and significantly higher than other methods, shown in Figure 3 (e-h). In contrast, MIFS is most sensitive to IR changes and most unstable. Moreover, when IR or the number of selected feature changes, CLAND is the least affected, which shows that our proposed method can stably obtain adequate information. Throughout the four evaluation criteria, CLAND showed the best accuracy and showed the best stability.

### 4.3 Biological significance of biomarker discovered by CLAND

Table 3 lists the top ten cancer genes obtained by CLAND for each cancer type. The genes are ranked by importance from high to low for distinguishing between cancer and normal samples. The completed results of all datasets are shown in Supplementary A.1. Besides classification performance and stability, we also consider the biological function of selected cancer genes. Take several genes in the table as illustrations. The gene with the highest score in KICH is *UMOD*, its *variants* are associated with chronic kidney disease in several studies [43], and its expression value is significantly down-regulated in renal cell carcinoma [44]. Because of the abundant expression in the colon but absence in colonic adenomas and adenocarcinomas, the *SLC26A3* is considered a potential tumor suppressor gene [45, 46]. The gene with the highest score in THCA is *TFF3*, which has been proved to be an oncogene in various types of cancers, such as breast, gastric and colorectal cancers [47, 48]. *TFF3* is a gene crucial in the signaling transduction pathway MAPK/ERK, which plays an essential role in tumor progression and metastasis, and can be used as a clinical therapeutic target for thyroid cancer [49]. *PER1*, with the highest score in BLCA, is a core in the generation of circadian rhythms, an essential regulator of cell division. The over-expression of *PER1* makes cancer cells sensitive to DNA damage-induced apoptosis, while the expression level of *PER1* in cancer patients is usually low [50]. Some other studies have shown that it plays a vital role in tumor occurrence, invasion, and prognosis[51].

**Table 3.**
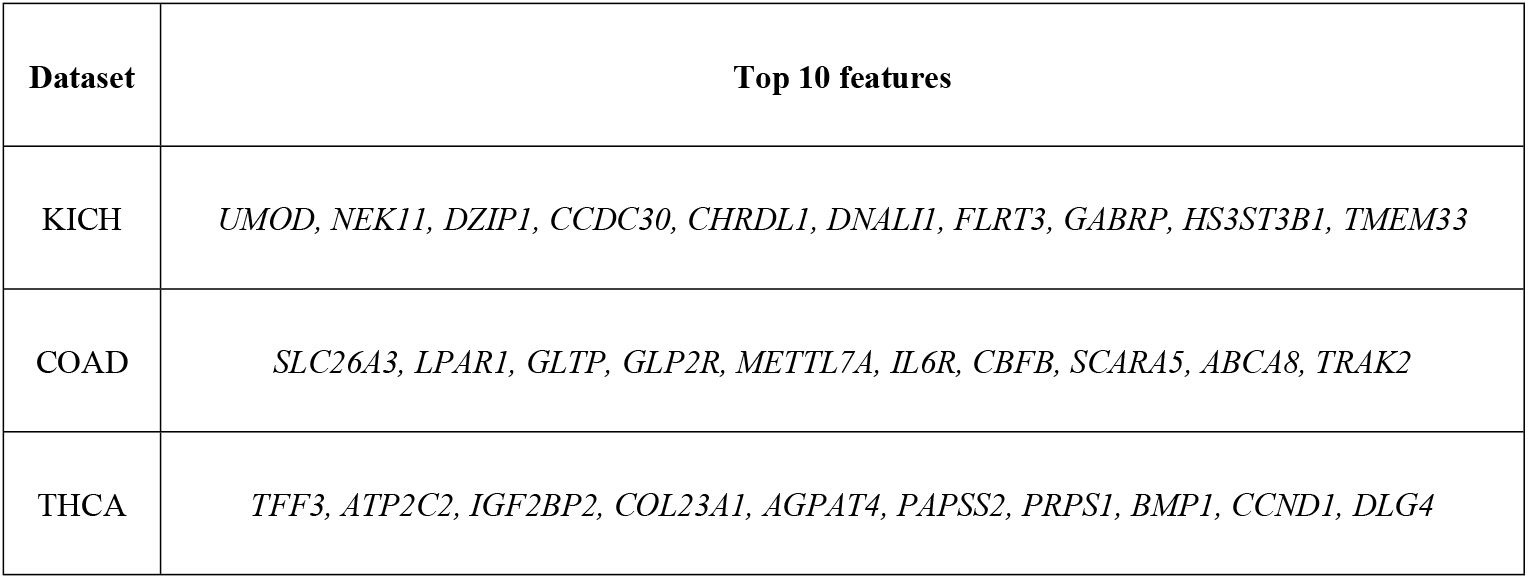
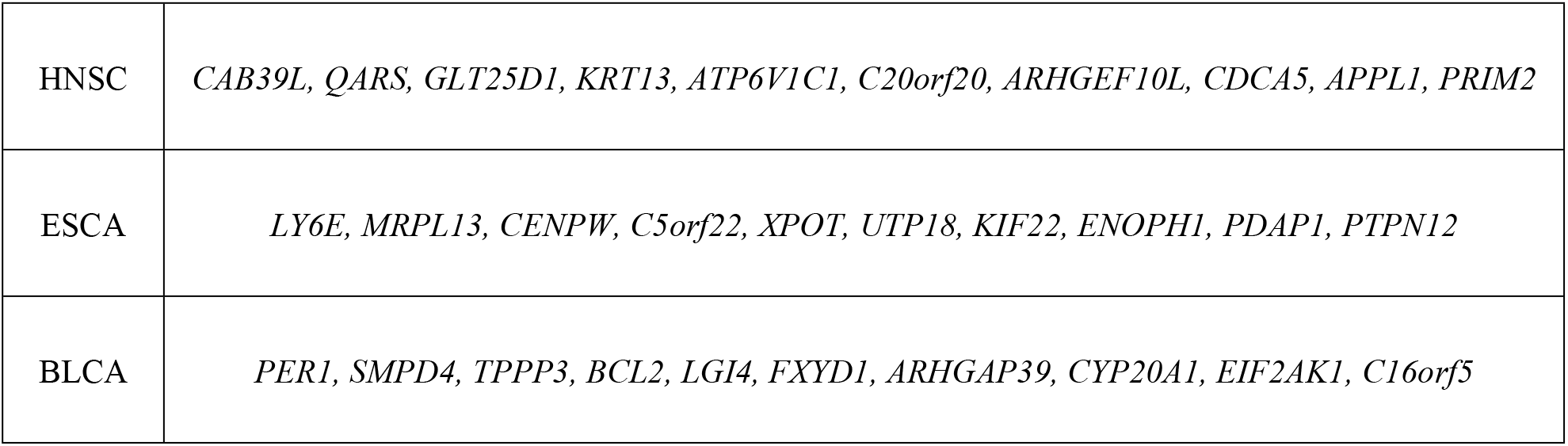
Biomarkers of cancers selected by CLAND

## Conclusion

This study describes how the class-imbalance problem affects the exploration of reliable biomarkers from cancer high-dimensional and non-linear omics data. By introducing the causal mechanism, we elucidate that class-imbalance reduces the stability of feature selection methods by simultaneously affecting the selected features and class prediction. Moreover, we propose a new feature selection method inspired by causality theory and technique called CLAND. We believe that the feature selection method should consider all the difficulties of biological datasets to obtain more valuable biomarkers. Therefore, the framework of CLAND is a dual-branch structure, including a class-wise causal branch and a sample-wise causal branch to eliminate the impact of imbalanced distribution. By conducting experiments on six representative real cancer data sets, CLAND has been proven to have better performance, better stability, broader applicability than state-of-the-art methods, and can find biomarkers with solid biological significance. In general, we provide a novel paradigm for feature selection from a causal perspective.

## Acknowledgments

The authors thank funding support from the National Natural Science Foundation of China (61902144, U19A2065, 61976102).

## Author Contribution

YL conceived and designed the model and performed the experiments. YL and QH analyzed the result. YL and HS wrote the paper. YC and HS carried out revision of the manuscript.

## Notes

### Competing Interest Statement

The authors have declared no competing interest.

